# Bacterial NDH-1 is a key gut target of metformin mediating its antidiabetic effects

**DOI:** 10.64898/2026.10.02.756213

**Authors:** Christina Isber, Linda Hadjadj, Lucile Pinault, Riyad Razzouk, Jean-Marc Rolain, Fadi Bittar

## Abstract

Metformin is the first-line therapy for type 2 diabetes, yet the mechanisms underlying its glucose-lowering effects remain incompletely understood. Increasing evidence implicates gut microbiota as an important mediator of metformin action, but the bacterial targets of the drug remain poorly defined. Here, we identify bacterial NADH oxidoreductase (NDH-1), the homolog of mitochondrial complex I, as a key target of metformin. Metformin inhibited bacterial growth at millimolar concentrations and, at subinhibitory concentrations, markedly impaired carbohydrate fermentation in *Escherichia coli* and other NDH-1-containing bacteria. Supplementation with nicotinic acid and Mg²⁺ restored fermentation, whereas pyruvate bypassed the metabolic defect, implicating impaired NAD⁺ regeneration and disruption of metabolism upstream of pyruvate formation. Genetic analysis further established NDH-1 dependence: metformin failed to inhibit fermentation in an *E. coli* ΔnuoF mutant lacking functional NDH-1 but retained activity in an NDH-2-deficient Δndh strain. Molecular docking predicted metformin binding within the NDH-1 quinone-binding region, proximal to conserved residues involved in quinone reduction. Consistent with this mechanism, *Staphylococcus aureus*, which lacks NDH-1, was not affected in carbohydrate fermentation, while several gut-associated probiotic and short-chain fatty acid-producing bacteria, including *Lactobacillus*, *Bifidobacterium*, and *Akkermansia muciniphila*, lacked detectable NDH-1 homologs, suggesting intrinsic resistance to this metabolic effect of metformin. These findings identify bacterial NDH-1 as a metabolic target of metformin and provide a mechanistic framework for its effects on the gut microbiota. Selective inhibition of NDH-1-dependent bacteria could alter microbial substrate utilization and ecological interactions in the intestine, potentially contributing to the microbiome-mediated component of metformin’s antidiabetic effects.

**Significance Statement:** Metformin is one of the most widely used drugs for type 2 diabetes, but how it interacts with the gut microbiota remains unclear. We show that metformin targets bacterial NADH oxidoreductase (NDH-1), the bacterial homolog of mitochondrial complex I, disrupting bacterial energy metabolism and carbohydrate fermentation. Bacteria lacking NDH-1, including gut-associated short-chain fatty acid-producing bacteria such as *Lactobacillus*, *Bifidobacterium*, and *Akkermansia muciniphila*, may therefore be less susceptible to this effect. Our findings identify a bacterial molecular target of metformin and provide a mechanistic framework linking drug-microbe interactions to microbiome remodeling. This work expands the understanding of metformin beyond host-directed mechanisms and highlights bacterial respiratory metabolism as a determinant of drug-microbiota interactions.

## Introduction

Non-antimicrobial drugs and their influence on microbiota have emerged as a growing concern. Many exhibit intrinsic antimicrobial activity, enabling them to inhibit bacterial growth (1), while promoting horizontal gene transfer (2), through conjugation (3) and natural transformation (4). Given that non-antimicrobial drugs constitute ∼95% of the global drug market (2), their widespread use, combined with environmental persistence, has led to their accumulation as pollutants at biologically relevant concentrations (5).

Metformin, the first-line treatment for type II diabetes mellitus (T2DM), represents a prominent example. It is among the most frequently detected active pharmaceutical ingredients in aquatic environments, including rivers, sediments, and wastewater systems (6–8). Originally derived from galegine, an active compound of *Galega officinalis* (9, 10), metformin was synthesized in 1922 (11) and introduced clinically in 1957 (12). Currently prescribed to more than 200 million individuals worldwide (13, 14), metformin has been associated with pleiotropic effects spanning metabolic, cardiovascular, oncological and infectious diseases (15–17).

Beyond its systemic effects on host metabolism, accumulating evidence indicates that metformin exerts a significant portion of its therapeutic action through the gut environment. Oral administration induces marked alterations in gut microbiota composition, including enrichment of taxa such as *Escherichia* spp. and *Akkermansia muciniphila* and depletion of others such as *Intestinibacter* spp (18). These compositional changes are functionally relevant, as they are associated with shifts in microbial metabolism, notably increased production of short-chain fatty acids (SCFAs), which contribute to improved glycemic control (19, 20). Consistent with this, the reduced efficacy of intravenous compared with oral administration (21) underscores the central role of the intestine and supports the contribution of microbiota-dependent mechanisms (19, 20). However, despite decades of clinical use, the molecular basis of metformin’s glucose-lowering effects remains incompletely resolved. Classical models emphasize hepatic actions, particularly suppression of gluconeogenesis, alongside inhibition of mitochondrial respiratory-chain complex I, which contributes to reduced blood glucose (19); yet these mechanisms do not fully account for its microbiome-mediated and gut-dependent effects. Importantly, microbiota alterations have also been implicated in metformin-associated gastrointestinal intolerance (22) and improved glycemic control (23, 24), highlighting the complexity of host–microbe–drug interactions.

As speculated, each bacterial protein, having a human protein homolog targeted by metformin, should be listed as a suspected target of metformin in bacteria (25). Given that NADH quinone oxidoreductase (NDH-1) is the functional and structural homolog of mitochondrial complex I, we hypothesize that metformin may target bacterial metabolic pathways via NDH-1 inhibition. In this study, we aimed to evaluate the antibacterial activity of metformin across a range of bacterial species through determination of minimum inhibitory concentrations (MICs), characterize its effects on bacterial physiological and metabolic profiles using standardized biochemical assays like Gallery API20E, and explore our hypothesis more through genomic and bioinformatic analyses. By integrating phenotypic, genetic and *in-silico* approaches, this work seeks to elucidate how metformin modulates bacterial systems and to establish a mechanistic link between its antimicrobial properties, microbiome alterations, and host metabolic outcomes.

## Materials and Methods

### Chemicals and stock solutions used

We used cation adjusted Mueller-Hinton broth media (CAMHB, Merck KGaA, Darmstadt, Germany), M9 minimal medium (Sigma Aldrich, St. Louis, MO, USA), Trypticase soy agar (TSA, bioMerieux, Marcy-l’Etoile, France), and MacConkey agar (TSA, bioMerieux, Marcy-l’Etoile, France) as mediums for bacterial culture. We used the following pharmaceuticals and chemicals: Metformin (Sigma Aldrich, Burlington MA, USA), Imipenem (Sigma Aldrich, Saint-Quentin-Fallavier, France), EDTA (ThermoFisher Scientific, Waltham, Massachusetts, USA), MgCl_2_ (Sigma Aldrich, St. Louis, MO, USA), Nicotinic acid (Sigma Aldrich, St. Louis, MO, USA), methyl Pyruvate (Sigma Aldrich, St. Louis, MO, USA), Glucose (Euromedex, Souffelweyersheim, France). For Imipenem, we prepared 1g/L stock solution in deionized water using a lyophilized formulation. Metformin was directly diluted to a concentration of 256 mg/ml in a sterile CAMHB. For the Modified API 20E system, we used the API20E strips and API STAPH strips (bioMerieux, Marcy-l’Etoile, France) containing dehydrated substrates.

### Bacterial strains and culture conditions

We analyzed bacterial strains belonging to ESKAPEE pathogenic strains, in addition to strains of *P. mendocina* isolated and characterized at the Institut Hospitalier Universitaire Méditerranée Infection (IHU-MI) (Marseille, France). All the strains used are listed in Table.1 along with their place of isolation, date of isolation and resistance profile. Three bacterial strains belonging to the Keio collection of *E. coli* K-12 were also used (BW25113, JW1095-1, JW2279-3) (26). All bacteria were cultured on TSA, MacConkey agar, CAMHB and M9 minimal medium for 24h of incubation at 37°C. Fresh bacterial growth was used for all analyses.

### Minimum inhibitory concentration (MIC) assays

MICs of Metformin and Imipenem were determined using the broth microdilution method. The sterile CAMHB medium was used as negative control, while the *E. coli* CIP7624 type strain, with known antibiotic sensibility, served as positive control. Visualization of bacterial growth was done by adding iodonitrotetrazolium to the wells. Same protocol was used with M9 as medium in the plates for *E. coli* strains.

### Checkerboard assay

To assess the chelating ability of metformin, Imipenem was tested in combination with Metformin using the checkerboard test, as previously described by Sopirala et al. (27). Serial dilutions of each drug at decreasing concentrations were prepared in CAMHB ranging from 4*MIC to 1/128*MIC. Each checkerboard well was streaked with 45 μL of each agent and 10 μL bacterial suspension at 10^5^ CFU/ml. Additional rows were used to determine the MIC of each used drug alone by adding 90 μL of each agent. The Fractional inhibitory concentration (FIC) and the Fractional inhibitory concentration index (FICI) were calculated according to the following equation FICI = FIC_M_ + FIC_A_ = (MIC of metformin in combination with imipenem/MIC of metformin alone) + (MIC of imipenem in combination with metformin/MIC of imipenem alone). The interaction was defined as synergistic (FICI ≤ 0.5), additive (FICI > 0.5 and < 1), indifferent (FICI >1 and < 2), and antagonistic (FICI > 2).

### Modified-API (Analytical Profile Index) 20E system

The commonly known protocol of the gallery API20E system was used with slight modification. A 0.5 McFarland bacterial inoculum was prepared with an overnight culture using 1.5% saline solution. For each bacterial strain tested, multiple strips were inoculated, including one control strip containing only the bacterial inoculum and additional strips containing bacterial inoculum supplemented with the different tested combinations of metformin, EDTA, MgCl₂, nicotinic acid, methyl pyruvate and glucose. The main concentration of metformin tested was half-MIC. For EDTA and MgCl_2_, a concentration of 2mM was tested. For Nicotinic acid, a series of concentrations were tested ranging between 200µM and 1mM. For methyl pyruvate, a concentration of 2mM was tested. For glucose, a concentration of 0.5M was used to provide excess glucose in each well. The final solutions, containing the combination of the different used compounds, were checked for 0.5 McFarland concentration required for gallery API. 100 µl of the bacterial solution were added to each well in the strip. Sterile paraffin oil was used to seal the wells requiring anaerobic conditions. The incubation temperature of the strip was maintained at 37°C with an incubation time of 24 hours. API STAPH strips were used to test the strains of *S. aureus* following the same protocol. The results were interpreted according to the guidelines of color change for a positive or negative reaction.

### Molecular Docking and Visualization

Molecular docking studies were conducted using AutoDock Vina (Version 1.2.7). The three-dimensional (3D) protein structure of NDH-1 complex (PDB ID: 7P63) was obtained from the RCSBD Protein Data Bank and prepared using AutoDock tools4 (Version 1.5.7) with polar hydrogen bonds added and Kollman charges assigned. Ligand restraints were obtained from PubChem database (Metformin: CID 4091; Fluorene CID 6853; Decylubiquinone (DCQ) CID 2971) and prepared using AutoDock tools4 with hydrogen bonds added and Gasteiger charges assigned. Docking calculations were carried out using a grid box centered at (102.03, 108.15, 144.55) Å, corresponding to DCQ binding site, with dimensions of 24 × 24 × 24 Å, encompassing the target binding region of the protein. The exhaustiveness parameter was set to 32, with a maximum of 10 binding modes generated per ligand within an energy range of 5 kcal/mol. The docking pose with the best Vina score was selected for further visualization and analyses. Protein-ligand interaction figures were generated using UCSF ChimeraX v1.11.1

### Genome acquisition

Complete genome sequences of three bacterial species (*Lactobacillus*, *Bifidobacterium*, and *Akkermansia muciniphila*) were retrieved from the National Center for Biotechnology Information (NCBI) Genome database. For each species, only assemblies annotated as “Complete Genome” were selected. In addition, the reference genome sequence of *E. coli* K-12 MG1655 was also retrieved from NCBI and annotated using prokka v1.14.6. The protein sequences, corresponding to the *Nuo* operon (the operon encoding for NDH-1, also known as NADH-quinone oxidoreductase) and consisting of 12 subunits, were extracted from the annotated reference genome. All genome datasets were accessed and downloaded in May 2026

### BLAST analysis for *Lactobacillus* species, *Bifidobacterium* species and *Akkermansia muciniphila* against reference *Nuo* operon

The search for sequence similarity was carried out using the BLAST (Basic Local Alignment Search Tool) algorithm. The *Nuo* operon sequence was queried against *Lactobacillus*, *Bifidobacterium*, and *Akkermansia muciniphila* database protein sequences using BLASTp. Alignments with a percent identity ≥ 80% and a query coverage ≥ 70% were considered significant and retained for further analysis.

### Ethics statement

This study was conducted exclusively using bacterial strains and did not involve human-derived samples or human participants. Consequently, the research did not fall within the scope of French regulations governing research involving human subjects, including Decree No. 2016-1537 of 16 November 2016 implementing Law No. 2012-300 of 5 March 2012. Therefore, ethical approval and informed consent were not applicable to this study.

## Results

### Minimum inhibitory concentration assay

For metformin, the MIC values of the tested species are listed in Table 1. Metformin exhibited an antibacterial activity against all tested strains with a value ranging between 16 and 64 mg/ml. An MIC of 32 and 64 mg/ml, using Muller Hinton broth, was obtained among the strains tested whether they are gram-positive or gram-negative. In M9 minimal medium, the MIC of the *E. coli* QA2806 strain showed to be 8 mg/ml. No significant difference was observed between the different species. Concerning the antibiotic resistance profile of the tested strains, no significance was noted regarding any correlation between the MIC value and a specific resistance profile.

**Table 1.**
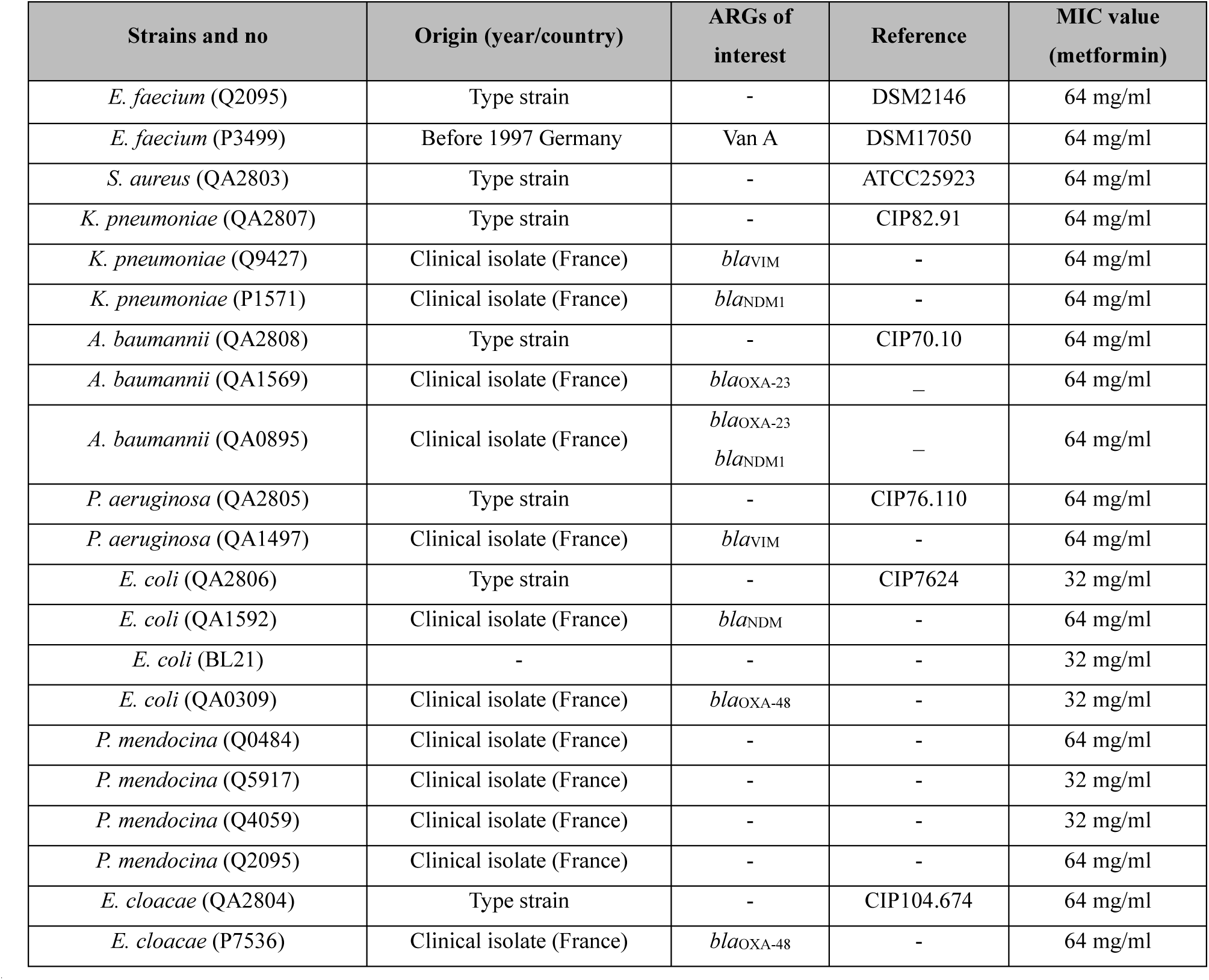
Details of strains used in this study with the minimal inhibitory concentration values for each strain obtained with Muller Hinton broth. ARGs: antibiotic resistance genes.

| Strains and no | Origin (year/country) | ARGs of interest | Reference | MIC value (metformin) |
| --- | --- | --- | --- | --- |
| *E. faecium* (Q2095) | Type strain | - | DSM2146 | 64 mg/ml |
| *E. faecium* (P3499) | Before 1997 Germany | Van A | DSM17050 | 64 mg/ml |
| *S. aureus* (QA2803) | Type strain | - | ATCC25923 | 64 mg/ml |
| *K. pneumoniae* (QA2807) | Type strain | - | CIP82.91 | 64 mg/ml |
| *K. pneumoniae* (Q9427) | Clinical isolate (France) | *bla*<sub>VIM</sub> | - | 64 mg/ml |
| *K. pneumoniae* (P1571) | Clinical isolate (France) | *bla*<sub>NDM1</sub> | - | 64 mg/ml |
| *A. baumannii* (QA2808) | Type strain | - | CIP70.10 | 64 mg/ml |
| *A. baumannii* (QA1569) | Clinical isolate (France) | *bla*<sub>OXA-23</sub> | — | 64 mg/ml |
| *A. baumannii* (QA0895) | Clinical isolate (France) | *bla*<sub>OXA-23</sub><br>*bla*<sub>NDM1</sub> | — | 64 mg/ml |
| *P. aeruginosa* (QA2805) | Type strain | - | CIP76.110 | 64 mg/ml |
| *P. aeruginosa* (QA1497) | Clinical isolate (France) | *bla*<sub>VIM</sub> | - | 64 mg/ml |
| *E. coli* (QA2806) | Type strain | - | CIP7624 | 32 mg/ml |
| *E. coli* (QA1592) | Clinical isolate (France) | *bla*<sub>NDM</sub> | - | 64 mg/ml |
| *E. coli* (BL21) | - | - | - | 32 mg/ml |
| *E. coli* (QA0309) | Clinical isolate (France) | *bla*<sub>OXA-48</sub> | - | 32 mg/ml |
| *P. mendocina* (Q0484) | Clinical isolate (France) | - | - | 64 mg/ml |
| *P. mendocina* (Q5917) | Clinical isolate (France) | - | - | 32 mg/ml |
| *P. mendocina* (Q4059) | Clinical isolate (France) | - | - | 32 mg/ml |
| *P. mendocina* (Q2095) | Clinical isolate (France) | - | - | 64 mg/ml |
| *E. cloacae* (QA2804) | Type strain | - | CIP104.674 | 64 mg/ml |
| *E. cloacae* (P7536) | Clinical isolate (France) | *bla*<sub>OXA-48</sub> | - | 64 mg/ml |

### The effect of the combination between metformin and imipenem

One review study described the ability of metformin to sequester metal ions such as Zn^2+^ (28). Based on this property and on the fact that carbapenemase enzymes (enzymes that degrade Carbapenem drug class) require Zn^2+^ to function, we thought of evaluating the effect of combining metformin with imipenem to determine whether any synergy and decrease in imipenem MIC would be observed. EDTA was included as a control. The experiment was conducted using one *K. pneumoniae* NDM-1 strain (P1571) and one *E. coli* NDM-5 strain (QA1592). The combination of metformin and imipenem showed an additive effect with an FIC of 0.75 for the *E. coli* tested strain and an FIC of 0.5625 for the *K. pneumoniae* tested strain. In contrast, EDTA, which served as the control, showed a synergistic effect with imipenem with an FIC index of 0.093 and 0.156 for *K. pneumoniae* and *E. coli* strains respectively.

### Metabolic pathways inhibition

Prepared *E. coli* inocula were incubated in API 20E strips with or without metformin, and metabolic profiles are summarized in Fig. 1. Treatment with metformin at ½ MIC resulted in complete inhibition of sugar fermentation across all tested strains. A concentration-dependent effect was observed, with full inhibition at ½ and ¼ MIC and partial inhibition at lower concentrations (Fig. 1A). For *S. aureus* strains, the addition of metformin at ½ MIC did not inhibit the acidification of glucose (Supplementary Fig 1). Similarly, EDTA (2 mM) abolished sugar fermentation, implicating metal ion dependency in fermentative metabolism. While MgCl₂ (2 mM) alone did not restore fermentation in metformin-treated samples, it fully reversed EDTA-mediated inhibition, indicating that the effect of EDTA is attributable to metal chelation (Fig. 1B). Nicotinic acid supplementation restored fermentation in a dose-dependent manner, with maximal effect at 1 mM, where partial recovery was observed in the presence of metformin. Notably, co-supplementation with nicotinic acid (1 mM) and MgCl₂ (2 mM) fully restored fermentation of all previously positive sugars despite metformin treatment. Furthermore, addition of methyl pyruvate (2 mM) in conjunction with metformin (½ MIC) resulted in positive sugar fermentation (Fig. 1C), suggesting that downstream metabolic intermediates can bypass the inhibitory effect.

**Fig 1.**
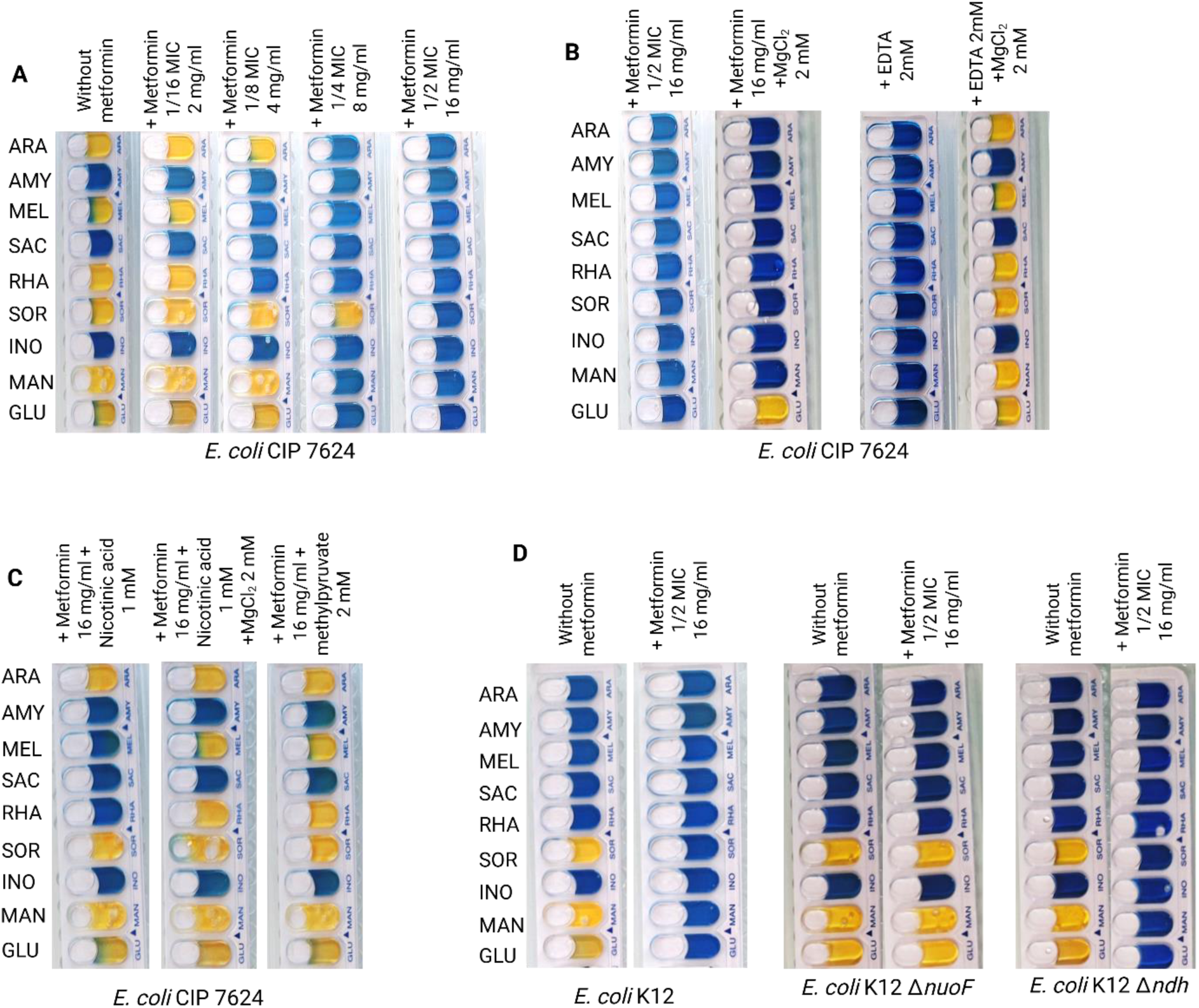
Metformin suppresses core carbohydrate fermentation pathways in *Escherichia coli*. **A,** Representative API 20E strips for *E. coli* CIP 7624 cultured in the absence and presence of increasing concentrations of metformin (1/16 to 1/2 MIC). Carbohydrate fermentation reactions display a progressive shift from yellow to blue with increasing metformin concentrations, reflecting a dose-dependent inhibition of carbohydrate fermentation. **B,** Addition of MgCl₂ to *E. coli* CIP 7624 cultured with metformin (1/2 MIC) did not restore carbohydrate fermentation, whereas supplementation with MgCl₂ directly within the API strip restored sugar fermentation following a confirmed EDTA-mediated inhibition. **C,** Addition of nicotinic acid (1 mM) partially restored carbohydrate fermentation, whereas its combination with MgCl₂ (2 mM) fully restored fermentation profiles. Supplementation with methyl pyruvate (2 mM) also led to positive fermentation reactions, indicating that cells retain the capacity to utilize pyruvate and sustain downstream fermentation processes. These results suggest that metformin inhibits carbohydrate fermentation upstream of pyruvate production within the glycolytic pathway. **D,** *E. coli* ΔnuoF mutant (NDH-1–deficient, relying on NDH-2) retained positive carbohydrate fermentation in the presence of metformin, whereas *E. coli* Δndh mutant (NDH-2–deficient, reliant on NDH-1) exhibited a loss of fermentation. These results indicate that metformin selectively impairs NDH-1–dependent respiration while sparing NDH-2, linking the inhibition of carbohydrate fermentation to disruption of the NDH-1 pathway.

To probe the mechanism of inhibition, *E. coli* Keio knockout strains (Δ*nuoF* and Δ*ndh*) were evaluated using the API 20E system in the presence and absence of metformin. In the Δ*nuoF* strain, lacking functional NDH-1, metformin did not inhibit sugar fermentation. In contrast, the Δ*ndh* strain, which lacks NDH-2 and relies on NDH-1, remained sensitive to metformin, exhibiting marked inhibition of fermentation (Fig. 1D). Together, these findings indicate that metformin selectively disrupts NDH-1–dependent metabolism, leading to impaired sugar fermentation. The integrity of the mutant strains was confirmed by a whole-genome sequencing, for the 2 mutant strains and the parental strain, where this sequencing verified the accurate disruption of the target genes and ruled out unintended mutations affecting other genes.

To test if excess glucose presence promotes the use of NDH-2, thus allowing sugar fermentation despite NDH-1 inhibition, additional glucose was supplied to the wells. Results showed that a concentration of 0.5M of additional glucose restored sugar fermentation in the presence of metformin in the *E. coli* QA2806 strain (Supplementary Fig. 2)

### Molecular Docking and Visualization

Molecular docking of metformin to the closed conformation of NDH-1 from *E. coli* was performed using AutoDock Vina. The top-ranked pose produced a Vina score of −4.9 kcal·mol⁻¹, indicating a moderate but stable interaction. Metformin binds within the quinone cavity at the interface of chains C/D, positioned above the quinone headgroup (DCQ) and adjacent to the N2 Fe–S cluster, consistent with a binding site at the base of the Q-channel (Fig. 2). Metformin interacts with key catalytic residues His228, Tyr322, and Asp592. Its biguanidinium core overlaps with the spatial region of the quinone headgroup, forming close contacts with Tyr322 and Asp592 while remaining proximal to His228. Structural and mechanistic studies indicate that quinone reduction in bacterial complex I involves a conserved catalytic network in subunit NuoCD (chains C/D), where histidine and tyrosine residues act as proton donors in conjunction with a conserved aspartate that undergoes protonation/deprotonation during quinone headgroup reduction (29). This placement introduces steric interference with the quinone-binding trajectory, constraining DCQ positioning within the pocket. Such interactions are likely to perturb the local hydrogen-bonding and electrostatic environment and interfere with the His228-mediated proton relay. Given the requirement for precise quinone alignment near the N2 cluster for efficient electron transfer, this binding mode supports a dual inhibitory mechanism involving restricted quinone access and disruption of the catalytic network.

**Fig 2.**
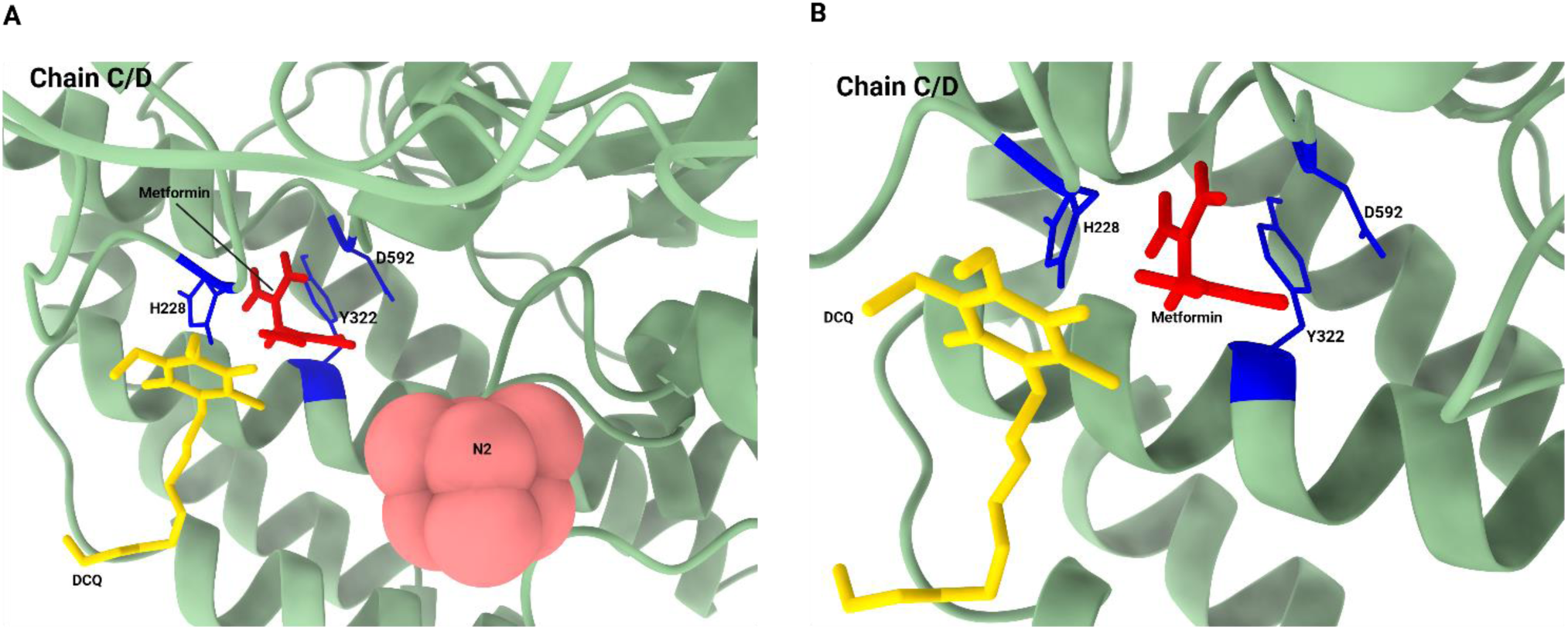
Docking predicts metformin occupation of the NDH-1 quinone-binding pocket. **A,** Metformin is positioned in proximity to key residues lining the quinone-binding pocket of NDH-1 (chains C/D shown in green ribbon representation), including histidine 228 (H228), tyrosine 322 (Y322), and aspartate 592 (D592) (blue), partially overlapping the binding region of the quinone analog DCQ (yellow). The nearby N2 iron–sulfur cluster (pink) highlights the spatial relationship between the predicted metformin binding site and the electron transfer pathway. **B,** Close-up view of the quinone-binding pocket highlighting the predicted metformin binding pose.

Docking protocol validation was performed using decylubiquinone (DCQ) as a positive control and fluorene, a decoy molecule, as a negative control consistent with standard docking validation approaches that employ non-binding molecules to assess specificity. Re-docking of DCQ reproduced its crystallographic binding pose within the quinone-binding site, with conserved interactions involving key residues (Supplementary Fig. 3A). In contrast, fluorene did not bind within this pocket and was instead located at a different site (chain H), with no interaction with quinone-associated residues (Supplementary Fig. 3B), supporting the specificity of the docking approach.

### In silico analysis for *Lactobacillus* species, *Bifidobacterium* species and *Akkermensia municiphila*

BLASTp analysis of the (*Escherichia coli* K-12 MG1655 *nuoA–nuoN* operon (GenBank accession: **NC_000913.3**)) against 3 different datasets of complete *Lactobacillus, Bifidobacterium, and Akkermensia municiphila* genomes retrieved from the NCBI Assembly database (assembly level: Complete Genome; accessed May 2026) databases returned with no significant hits with a cutoff of 80% for both the coverage and similarity. These results where the same for the three databases of *Lactobacilli, Bifidobacterium* and *Akkermansia muciniphila*.

## Discussion

The widespread presence of non-antimicrobial drugs in the environment highlights the continuous exposure of microbiota to these compounds and underscores the importance of understanding their bidirectional interactions with microbes. Metformin, one of the most widely used active pharmaceutical ingredients, can be degraded by bacteria via the *mfmAB* gene cluster (30, 31), with a recent capture by a plasmid in defined bacterial species reported (32), suggesting an adaptive microbial response to this compound. This interaction raises the possibility of intrinsic antimicrobial activity, which has so far been reported only in limited bacterial strains (33, 34). In this study, we demonstrate that metformin exerts antimicrobial activity across a broader range of species and propose a mechanism in which it inhibits bacterial sugar fermentation, particularly in *E. coli*. This effect may represent an initial step linking metformin’s microbial activity to its glucose-lowering action.

The antimicrobial activity of metformin suggests biologically relevant effects on a wide range of bacteria, including clinically important pathogens. In clinical practice, metformin is administered at 1–2 g daily (34) and accumulates in the intestinal lumen at concentrations 30–300-fold higher than in plasma (36), due to its slow and incomplete gastrointestinal absorption (37). Modeling studies estimate duodenal concentrations of 50–75 mM (8-12 mg/ml) prior to absorption (38), indicating that local intestinal levels are sufficient to exert antimicrobial or metabolic effects on resident microbiota. The observed inhibition of metabolic activity at sub-inhibitory concentrations of metformin strengthens the hypothesis that this drug primarily targets bacterial metabolism rather than exerting a purely bactericidal effect.

In the API 20E system, metformin primarily disrupted carbohydrate fermentation, indicating a broad effect on central bacterial metabolism. Because diverse sugars converge into the glycolytic pathway, this pattern supports targeting of a common metabolic node rather than substrate-specific processes (Fig. 3). Pathway reconstruction (39, 40) underscores the reliance of key *E. coli* glycolytic enzymes, particularly kinases, on Mg²⁺ as an essential cofactor. In light of the reported metal-chelating properties of metformin (28, 41), sequestration of Mg²⁺ represents a plausible mechanism underlying this metabolic inhibition. This interpretation is consistent with clinical observations linking metformin treatment to reduced circulating Mg²⁺ levels (42, 43), suggesting interference with magnesium homeostasis. However, the incomplete rescue of fermentation upon MgCl₂ supplementation, despite EDTA-induced inhibition, indicates that Mg²⁺ chelation alone does not fully account for the observed effects. Collectively, these data support a model in which Mg²⁺ limitation contributes to, but does not entirely explain, the inhibitory impact of metformin on bacterial glycolysis.

**Fig 3.**
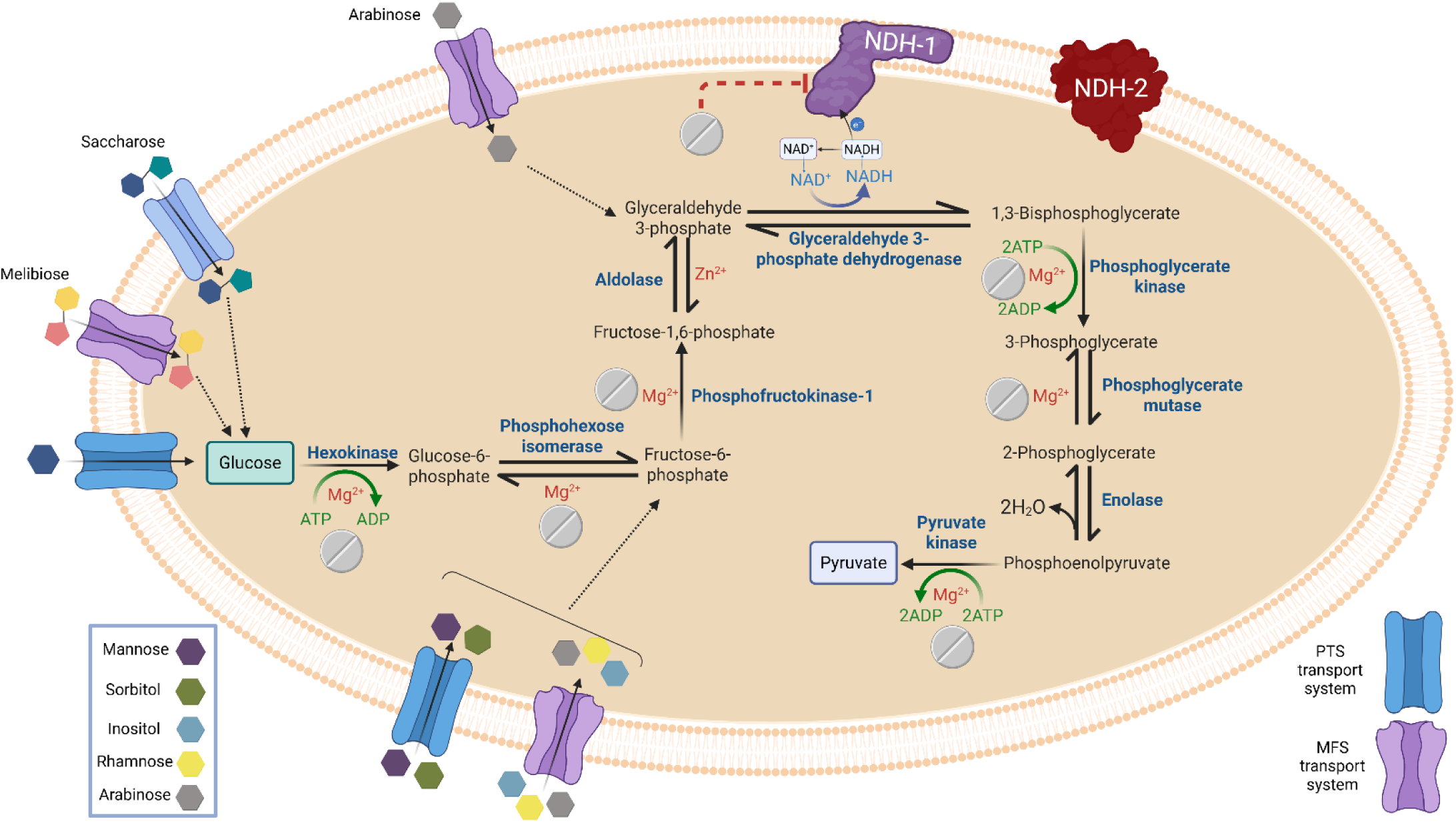
Proposed mechanism linking NDH-1 inhibition and Mg²⁺ chelation by metformin to its induced carbohydrate metabolic disruption in *Escherichia coli*. Schematic representation of metformin-induced perturbations of central metabolism in *E. coli*. Metformin is proposed to inhibit the respiratory complex NDH-1 at the membrane, impairing NADH oxidation and altering the cellular redox balance (NAD⁺/NADH). This effect impacts the activity of glyceraldehyde-3-phosphate dehydrogenase that requires NAD^+^ to function, limiting progression through glycolysis. In parallel, metformin-mediated chelation of Mg²⁺ interferes with multiple Mg²⁺-dependent enzymes of the glycolytic pathway, thereby reducing flux through carbohydrate metabolism. Consequently, pyruvate production is diminished, leading to impaired fermentation capacity. NDH-2 is shown as an alternative NADH dehydrogenase that is not inhibited.

Metformin is known to target mitochondrial complex I in eukaryotic cells, contributing to its glucose-lowering effect (22, 44, 45). A similar mechanism is occurring in bacteria via inhibition of NDH-1, the bacterial homolog of complex I (46). Such inhibition will impair NAD⁺ regeneration, limiting the activity of NAD⁺-dependent enzymes and disrupting glycolysis. Consistent with this hypothesis, restoration of sugar fermentation upon supplementation with nicotinic acid (vitamin B3), a precursor of NAD⁺, indicates that NAD⁺ availability is a key limiting factor under metformin treatment. The requirement for combined supplementation with nicotinic acid and MgCl₂ to achieve full recovery further suggests that both NAD⁺ depletion and Mg²⁺ limitation contribute to the metabolic effects of metformin. Together, these observations support NDH-1 inhibition as a central component of metformin’s mechanism of action in bacteria, with Mg²⁺ chelation playing a secondary role (Fig. 3). The relatively weak chelating capacity of metformin is further indicated by checkerboard assays showing additive rather than synergistic effects compared to the strong chelator EDTA. Although metformin has been reported to sequester Zn²⁺ (28), the limited reduction in imipenem MIC (one to two dilutions) suggests that its zinc-chelating activity is insufficient to significantly inhibit carbapenemase function. Collectively, these observations indicate that metal chelation contributes to, but does not dominate, the antibacterial mechanism of metformin. Moreover, the effect of nicotinic acid underscores the importance of NAD⁺ availability.

Supplementation with methyl pyruvate further supports this mechanism, as bacteria maintained lactic acid production in the presence of metformin, indicating that downstream pyruvate utilization remains intact. This implicates that metformin inhibits a step upstream of pyruvate formation within glycolysis (Fig. 3).

Consistent with previous studies showing that NAD⁺ availability modulates sensitivity to metformin (47, 48), the higher MIC observed in Muller Hinton broth (32 mg/ml) compared to M9 minimal medium (8 mg/ml) can be explained by differences in nutrient composition. Muller Hinton broth contains vitamin B3 precursors and amino acids that support NAD⁺ synthesis, thereby mitigating metformin’s effect, whereas nutrient limitation in M9 medium forces *E. coli* to rely on glycolysis and fermentation (49, 50), increasing susceptibility. These observations further support the role of NAD⁺ depletion in metformin’s mechanism of action (Fig. 3).

The recent identification of *E. coli* agmatinase as a metformin target (35), together with our demonstration of NDH-1 inhibition, supports a model in which metformin directly modulates multiple bacterial metabolic pathways. Although agmatinase inhibition was proposed to result from competitive binding, the Mn^2+^-metal dependency of the enzyme raises the possibility that metal coordination may also contribute, consistent with our observations of mild Mg²⁺ chelation by metformin.

Further evidence supporting NDH-1 as a key metabolic target of metformin is provided by mutant analysis where results indicate that metformin selectively targets the NDH-1 complex, while sparing NDH-2. This specificity highlights NDH-1 as a critical component required for maintaining glycolytic activity under metformin exposure (Fig. 3).

Furthermore, the failure of metformin to inhibit sugar fermentation in *S. aureus* further supports NDH-1 as a key bacterial target of the drug, given that this species encodes only NDH-2 and lacks the NDH-1 complex (51).

The NDH-1 complex in bacteria consists of 12 subunits, with the C/D subunit harboring a conserved His–Tyr–Asp triad that is central to the proton-coupled electron transfer network responsible for quinone stabilization and proton delivery during reduction (29, 52). Because efficient electron transfer in complex I depends on the tight coupling between quinone positioning and proton relay, this catalytic region is particularly sensitive to perturbation, where even subtle disruptions can significantly reduce turnover. The predicted positioning of metformin within this environment suggests that its presence could interfere with the functional integrity of this triad, thereby affecting quinone chemistry rather than acting through direct channel occlusion. This interpretation is consistent with the model proposed by He et al. (2025), in which metformin behaves as a conformation-dependent inhibitor that becomes trapped near the ubiquinone redox site, within the same catalytic environment, without fully blocking substrate access (53).

Previous studies have shown that metformin induces a shift in the gut microbiota, characterized by a reduction in pro-inflammatory bacteria and an increase in beneficial taxa, contributing to its glucose-lowering effects (19, 20). In line with these observations, our findings suggest that metformin-mediated inhibition of bacterial sugar fermentation may selectively suppress the metabolic activity of organisms such as *Escherichia coli*. This effect could increase the availability of fermentable substrates within the intestinal lumen, thereby promoting the growth and activity of beneficial bacteria, including *Lactobacillus*, *Bifidobacterium*, and *Akkermansia muciniphila*, which are known producers of short-chain fatty acids (SCFAs). These metabolites play a central role in glucose homeostasis by stimulating GLP-1 secretion, enhancing insulin sensitivity, and reducing inflammation (19, 20) (Fig. 4).

**Fig 4.**
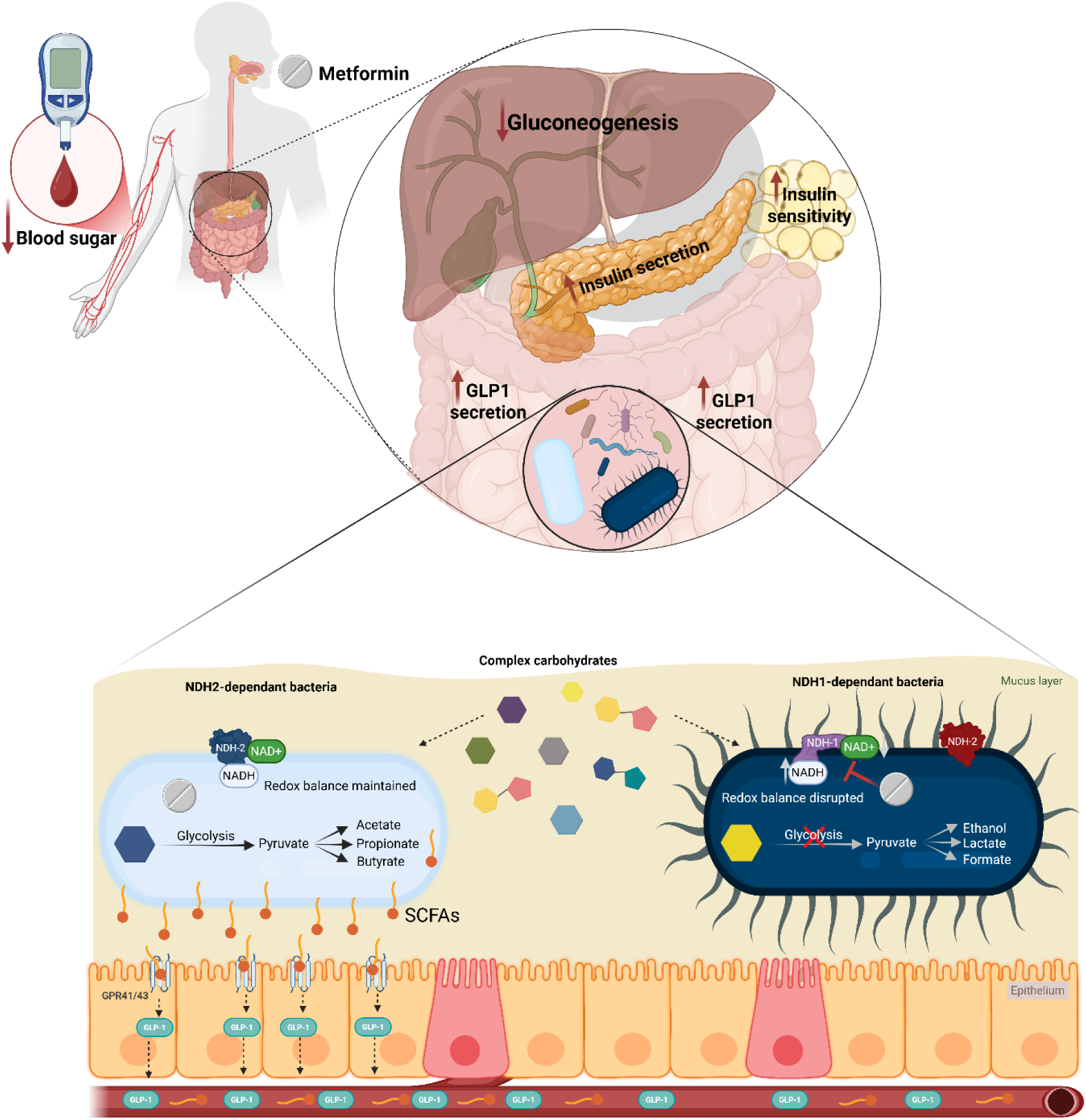
Proposed microbiome-mediated mechanism underlying the metabolic effects of metformin. Schematic representation of the impact of metformin on gut microbial composition and host glucose metabolism. Metformin selectively inhibits carbohydrate metabolism in NDH-1–dependent bacteria by targeting NDH-1, leading to disruption of redox balance (NAD⁺/NADH) and reduced glycolytic flux in these populations. In contrast, NDH-2–dependent bacteria maintain redox homeostasis, as NDH-2 is not inhibited by metformin, allowing continued carbohydrate metabolism. As a result, carbohydrate substrates become more available to NDH-2–dependent bacteria, which convert them into short-chain fatty acids (SCFAs), including acetate, propionate, and butyrate. Increased SCFA production stimulates glucagon-like peptide-1 (GLP-1) secretion from intestinal epithelial cells via G protein–coupled receptors (e.g., GPR41/43). Elevated GLP-1 levels enhance pancreatic insulin secretion and improve peripheral insulin sensitivity, while concurrently suppressing hepatic gluconeogenesis. Together, these microbiome-driven processes contribute to reduced blood glucose levels, supporting a model in which the therapeutic effects of metformin are partly mediated through modulation of gut microbial metabolism.

Importantly, the absence of the NDH-1 complex, identified here as a key target of metformin, in these bacterial taxa, as indicated by BLASTp analysis, suggests intrinsic resistance to metformin’s inhibitory effects on carbohydrate metabolism (Fig. 3 and Fig. 4). Thus, selective targeting of NDH-1–dependent bacteria may represent a mechanistic link between metformin’s antimicrobial activity, microbiota remodeling, and its systemic metabolic benefits.

Moreover, the observed recovery of sugar fermentation under excess glucose conditions and the presence of a functional NDH-2 suggest that increased glucose flux promotes a metabolic shift toward NDH-2 mediated respiration. A study by G. N. Vemuri et al. reported that high glucose flux is associated with increased expression of the *ndh* gene, resulting in higher levels of NDH-2 (54). Under high glucose conditions, enhanced glycolytic activity produces higher amounts of NADH and increases the demand for quicker NADH oxidation to maintain the redox balance. NDH-2 plays an important role in this quick NADH oxidation because, unlike NDH-1, NDH-2 is not coupled to proton translocation (55). Consequently, increased NDH-2 activity may sustain elevated glucose catabolism, reducing the amount of glucose available for absorption into the bloodstream and contributing to lower blood glucose levels.

## Conclusion

Gut science has been the hottest area of biomedicine for more than a decade, with gut microbiota emerging as the missing part of the puzzle in understanding the pathogenesis and treatments of many diseases. In this study, we uncovered the ability of metformin to inhibit the fermentation of sugars in *E. coli* by inhibiting the NDH-1 complex. This inhibition will favor the use of the available glucose and other sugars by other SCFAs producing bacteria, whose metabolites have been implicated in improved glucose homeostasis and insulin sensitivity.

Beyond its mechanistic implications, our findings highlight several potential determinants of bacterial resistance to metformin. First, enzymatic resistance mediated by the recently described *mfmAB* genes and their potential horizontal transfer may promote the dissemination of metformin-degrading capabilities within microbial communities. Second, intrinsic resistance appears to be closely associated with the absence of the drug target NDH-1, as illustrated by probiotic lactic acid bacteria, *Staphylococcus aureus*, and the *E. coli* K-12 Δ*nuoF* mutant. These observations further support the central role of NDH-1 in bacterial susceptibility to metformin. Finally, future studies should determine whether naturally occurring or adaptive mutations affecting NDH-1 structure and function constitute an additional mechanism of acquired resistance and contribute to the heterogeneous clinical response to metformin observed among patients.

More broadly, the identification of NDH-1 as a bacterial target of metformin highlights respiratory metabolism as a promising therapeutic vulnerability and supports the exploration of selective NDH-1 inhibitors as a new class of microbiome-modulating agents for metabolic and other chronic diseases, given the growing evidence of metformin’s clinical benefits in cancer and metabolic diseases (56) such as nonalcoholic steatohepatitis (NASH) (57).

## Ethical approval

not applicable

## Declaration of generative AI and AI-assisted technologies in the manuscript preparation process

During the preparation of this work the authors used ChatGPT to improve the grammar of some sentences. After using this service, the authors reviewed and edited the content as needed and took full responsibility for the content of the published article.

---

**Supplementary Fig 1.**
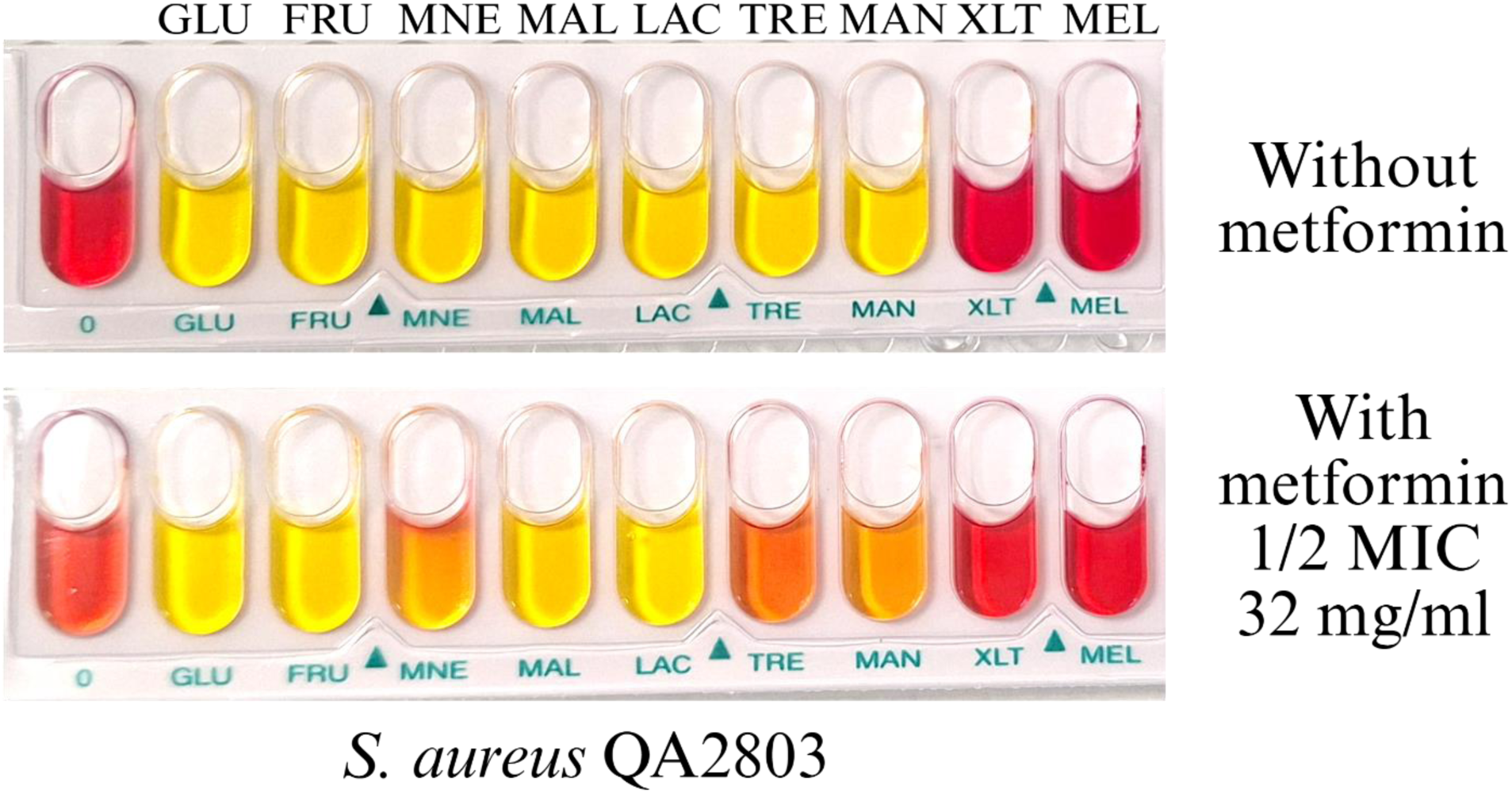
Metformin failed inhibition of carbohydrates acidification in *S. aureus*. Representative API STAPH strips for *S. aureus* QA2803 cultured in the absence and presence of metformin (1/2 MIC). Carbohydrate acidification reactions display a shift from red to yellow even in the presence of metformin, reflecting a lack of detectable inhibition of carbohydrate acidification. This observation is consistent with the lack of NDH-1 and the reliance of *S. aureus* species on NDH-2.

**Supplementary Fig 2.**
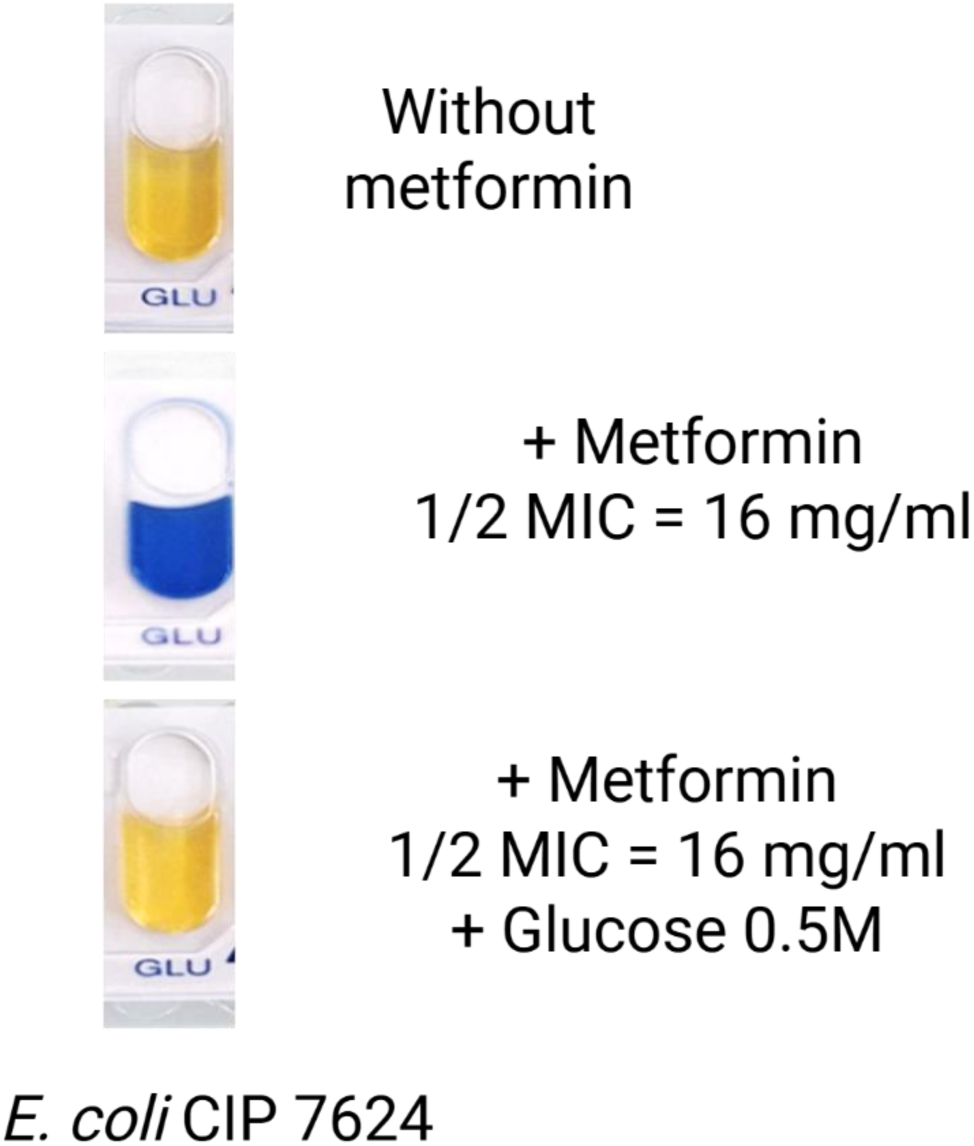
Excess glucose restored glucose fermentation in the presence of Metformin. Representative API 20E strips for *E. coli* QA2806 cultured in the presence and absence of metformin with a third strip characterized by the addition of 0.5M concentration of glucose. Glucose fermentation reactions display a shift in color from blue to yellow in the presence of both metformin and glucose reflecting a compensation of metformin inhibition in the presence of excess glucose.

**Supplementary Fig 3.**
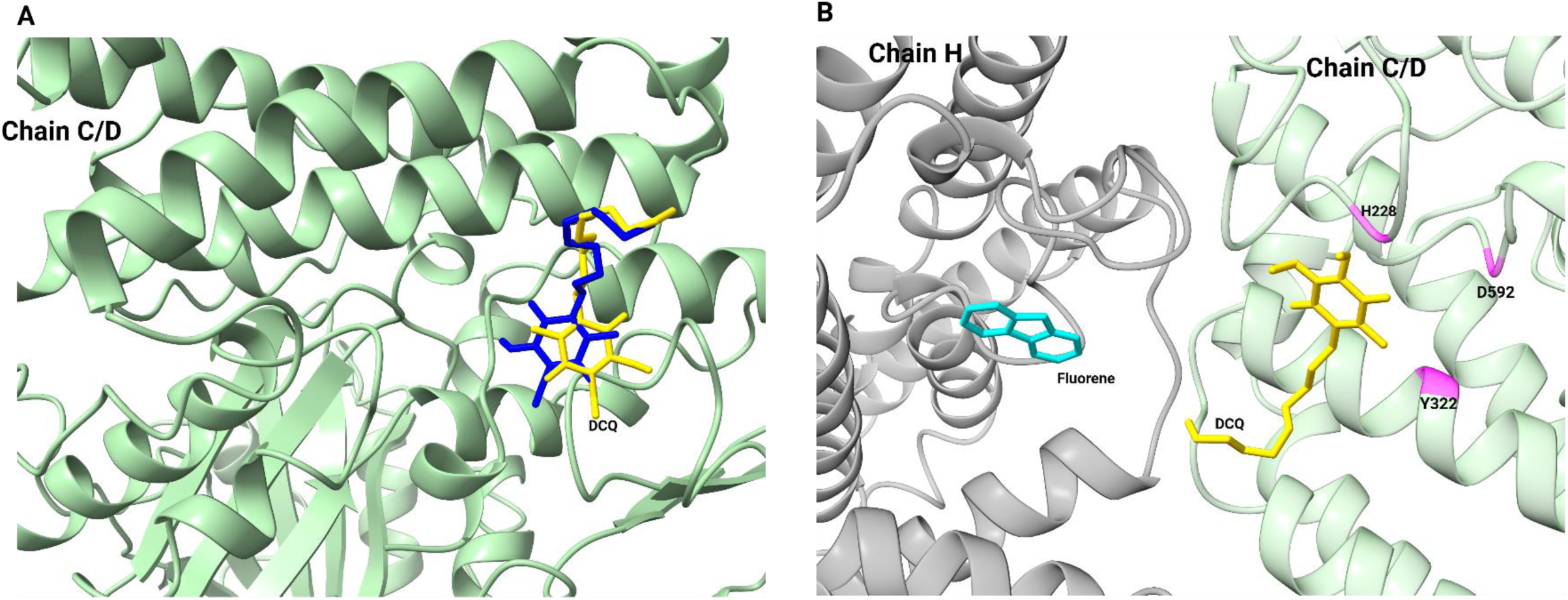
Validation of the docking protocol using a positive and a negative control. **A,** re-docking of DCQ (positive control) showed correct positioning of the DCQ in the quinone-binding site and reproduced its binding pose. **B,** Fluorene (negative control), showed a different binding pose in chain H with no interaction with key quinone-binindg site residues. These results supports the reliability and specificity of the docking results.

## References

1. L. Maier, et al., Extensive impact of non-antibiotic drugs on human gut bacteria. Nature 555, 623–628 (2018).

2. A. Hayes, et al., Common non-antibiotic drugs enhance selection for antimicrobial resistance in mixture with ciprofloxacin. [Preprint] (2025). Available at: http://biorxiv.org/lookup/doi/10.1101/2025.03.23.644574 [Accessed 26 June 2026].

3. Y. Wang, et al., Non-antibiotic pharmaceuticals promote the transmission of multidrug resistance plasmids through intra- and intergenera conjugation. The ISME Journal 15, 2493–2508 (2021).

4. Y. Wang, et al., Non-antibiotic pharmaceuticals enhance the transmission of exogenous antibiotic resistance genes through bacterial transformation. The ISME Journal 14, 2179–2196 (2020).

5. N. Aryal, et al., Fate of environmental pollutants: A review. Water Environment Research 92, 1587–1594 (2020).

6. J. L. Wilkinson, et al., Pharmaceutical pollution of the world’s rivers. Proc. Natl. Acad. Sci. U.S.A. 119, e2113947119 (2022).

7. C. Littlejohn, et al., Environmental Concentrations of the Type 2 Diabetes Medication Metformin and Its Transformation Product Guanylurea in Surface Water and Sediment in Ontario and Quebec, Canada. Environmental Toxicology and Chemistry 42, 1709–1720 (2023).

8. R. M. Briones, W.-Q. Zhuang, A. K. Sarmah, Biodegradation of metformin and guanylurea by aerobic cultures enriched from sludge. Environmental Pollution 243, 255–262 (2018).

9. C. J. Bailey, Metformin: historical overview. Diabetologia 60, 1566–1576 (2017).

10. C. Bailey, C. Day, Metformin: its botanical background. Pract Diab Int 21, 115–117 (2004).

11. C. R. Triggle, et al., Metformin: Is it a drug for all reasons and diseases? Metabolism 133, 155223 (2022).

12. E. Alphonsewerner. Jamebesll, CCXIV.-The Preparation of Methylguanidine, and of Pp-Dimethylguanidine by the Interaction of Di-cyanodiamide,and Methylammonium and Dimethyl-ammonium Chlorides Respectively.

13. G. Schernthaner, G.-H. Schernthaner, The right place for metformin today. Diabetes Research and Clinical Practice 159, 107946 (2020).

14. E. Ahmad, et al., Where Does Metformin Stand in Modern Day Management of Type 2 Diabetes? Pharmaceuticals 13, 427 (2020).

15. I. H. Vieira, L. M. Barros, C. F. Baptista, D. M. Rodrigues, I. M. Paiva, Recommendations for Practical Use of Metformin, a Central Pharmacological Therapy in Type 2 Diabetes. Clinical Diabetes 40, 97–107 (2022).

16. S. Mohamed, Metformin: Diverse molecular mechanisms, gastrointestinal effects and overcoming intolerance in type 2 Diabetes Mellitus: A review. Medicine 103, e40221 (2024).

17. L. Zhu, et al., Association of Blood Glucose Control and Outcomes in Patients with COVID-19 and Pre-existing Type 2 Diabetes. Cell Metabolism 31, 1068–1077.e3 (2020).

18. T. Bryrup, et al., Metformin-induced changes of the gut microbiota in healthy young men: results of a non-blinded, one-armed intervention study. Diabetologia 62, 1024–1035 (2019).

19. C. B. Lee, S. U. Chae, S. J. Jo, U. M. Jerng, S. K. Bae, The Relationship between the Gut Microbiome and Metformin as a Key for Treating Type 2 Diabetes Mellitus. IJMS 22, 3566 (2021).

20. I. Szymczak-Pajor, J. Drzewoski, M. Kozłowska, J. Krekora, A. Śliwińska, The Gut Microbiota-Related Antihyperglycemic Effect of Metformin. Pharmaceuticals 18, 55 (2025).

21. J. Drewe, M. Foretz, S. Krähenbühl, Metformin—mechanisms of its glycemia-reducing effect. Pharmacological Reviews 78, 100106 (2026).

22. C. R. Reczek, et al., Metformin targets mitochondrial complex I to lower blood glucose levels. Sci. Adv. 10, eads5466 (2024).

23. E. Amengual-Cladera, et al., Metformin: From Diabetes to Cancer—Unveiling Molecular Mechanisms and Therapeutic Strategies. Biology 13, 302 (2024).

24. H. Wu, et al., Metformin modulates the gut microbiome in a mice model of high-fat diet-induced glycolipid metabolism disorder. BMJ Open Diab Res Care 10, e003149 (2022).

25. P. V. Bauer, et al., Metformin Alters Upper Small Intestinal Microbiota that Impact a Glucose-SGLT1-Sensing Glucoregulatory Pathway. Cell Metabolism 27, 101–117.e5 (2018).

26. Baba T, Ara T, Hasegawa M, Takai Y, Okumura Y, Baba M, et al. Construction of *Escherichia coli* K-12 in-frame, single-gene knockout mutants: the Keio collection. Mol Syst Biol. 2006;2: 2006.0008. doi:10.1038/msb4100050

27. M. M. Sopirala, et al., Synergy Testing by Etest, Microdilution Checkerboard, and Time-Kill Methods for Pan-Drug-Resistant Acinetobacter baumannii. Antimicrobial Agents and Chemotherapy 54, 4678–4683 (2010).

28. D. A. Rusanov, J. Zou, M. V. Babak, Biological Properties of Transition Metal Complexes with Metformin and Its Analogues. Pharmaceuticals 15, 453 (2022).

29. C. Harter, et al., Quinone chemistry in respiratory complex I involves protonation of a conserved aspartic acid residue. FEBS Letters 598, 2856–2865 (2024).

30. B. M. Martinez-Vaz, et al., Wastewater bacteria remediating the pharmaceutical metformin: Genomes, plasmids and products. Front. Bioeng. Biotechnol. 10, 1086261 (2022).

31. T. Li, et al., Discovery of a Ni2+-dependent heterohexameric metformin hydrolase. Nat Commun 15, 6121 (2024).

32. T. Vo, et al., Global lessons from antibiotic resistance: Metformin-hydrolysing genes in transposable elements, a new threat for type II diabetic patients? Journal of Global Antimicrobial Resistance 51, 29–34 (2026).

33. M. M. Masadeh, K. H. Alzoubi, M. M. Masadeh, Z. O. Aburashed, Metformin as a Potential Adjuvant Antimicrobial Agent Against Multidrug Resistant Bacteria. CPAA Volume 13, 83–90 (2021).

34. S. M. Shafik, H. A. Abbas, N. Yousef, M. M. Saleh, Crippling of Klebsiella pneumoniae virulence by metformin, N-acetylcysteine and secnidazole. BMC Microbiol 23, 229 (2023).

35. L. J. Tassoulas, L. P. Wackett, Insights into the action of the pharmaceutical metformin: Targeted inhibition of the gut microbial enzyme agmatinase. iScience 27, 108900 (2024).

36. L. J. McCreight, C. J. Bailey, E. R. Pearson, Metformin and the gastrointestinal tract. Diabetologia 59, 426–435 (2016).

37. E. Sundelin, J. B. Jensen, S. Jakobsen, L. C. Gormsen, N. Jessen, Metformin Biodistribution: A Key to Mechanisms of Action? The Journal of Clinical Endocrinology & Metabolism 105, 3374–3383 (2020).

38. C. J. Kelly, et al., Metformin inhibits digestive proteases and impairs protein digestion in mice. Journal of Biological Chemistry 299, 105363 (2023).

39. M. Kanehisa, S. Goto, KEGG: Kyoto Encyclopedia of Genes and Genomes. Nucleic Acids Res 28, 27–30 (2000).

40. N. Grüning, et al., The return of metabolism: biochemistry and physiology of glycolysis. Biological Reviews 101, 751–803 (2026).

41. L. Logie, et al., Cellular Responses to the Metal-Binding Properties of Metformin. Diabetes 61, 1423–1433 (2012).

42. F. Waanders, et al., Hypomagnesaemia and its determinants in a contemporary primary care cohort of persons with type 2 diabetes. Endocrine 67, 80–86 (2020).

43. K. E. Peters, S. A. P. Chubb, W. A. Davis, T. M. E. Davis, The Relationship between Hypomagnesemia, Metformin Therapy and Cardiovascular Disease Complicating Type 2 Diabetes: The Fremantle Diabetes Study. PLoS ONE 8, e74355 (2013).

44. E. Fontaine, Metformin-Induced Mitochondrial Complex I Inhibition: Facts, Uncertainties, and Consequences. Front. Endocrinol. 9, 753 (2018).

45. S. Martín-Rodríguez, P. De Pablos-Velasco, J. A. L. Calbet, Mitochondrial Complex I Inhibition by Metformin: Drug–Exercise Interactions. Trends in Endocrinology & Metabolism 31, 269–271 (2020).

46. R. Vashisht, S. K. Brahmachari, Metformin as a potential combination therapy with existing front-line antibiotics for Tuberculosis. J Transl Med 13, 83 (2015).

47. M. Parisotto, et al., The NAMPT Inhibitor FK866 Increases Metformin Sensitivity in Pancreatic Cancer Cells. Cancers 14, 5597 (2022).

48. D. Y. Gui, et al., Environment Dictates Dependence on Mitochondrial Complex I for NAD+ and Aspartate Production and Determines Cancer Cell Sensitivity to Metformin. Cell Metabolism 24, 716–727 (2016).

49. B. J. Koebmann, H. V. Westerhoff, J. L. Snoep, D. Nilsson, P. R. Jensen, The Glycolytic Flux in *Escherichia coli* Is Controlled by the Demand for ATP. J Bacteriol 184, 3909–3916 (2002).

50. N. Okahashi, S. Kajihata, C. Furusawa, H. Shimizu, Reliable Metabolic Flux Estimation in Escherichia coli Central Carbon Metabolism Using Intracellular Free Amino Acids. Metabolites 4, 408–420 (2014).

51. The two alternative NADH:quinone oxidoreductases from Staphylococcus aureus: two players with different molecular and cellular roles | Microbiology Spectrum. Available at: https://journals.asm.org/doi/full/10.1128/spectrum.04152-23 [Accessed 15 September 2026].

52. A. P. Gamiz-Hernandez, A. Jussupow, M. P. Johansson, V. R. I. Kaila, Terminal Electron– Proton Transfer Dynamics in the Quinone Reduction of Respiratory Complex I. J. Am. Chem. Soc. 139, 16282–16288 (2017).

53. Z. He, et al., Hydrophilic metformin and hydrophobic biguanides inhibit mitochondrial complex I by distinct mechanisms. Nat Struct Mol Biol 33, 100–111 (2026).

54. G. N. Vemuri, E. Altman, D. P. Sangurdekar, A. B. Khodursky, M. A. Eiteman, Overflow Metabolism in Escherichia coli during Steady-State Growth: Transcriptional Regulation and Effect of the Redox Ratio. Appl Environ Microbiol 72, 3653–3661 (2006).

55. A. M. P. Melo, T. M. Bandeiras, M. Teixeira, New Insights into Type II NAD(P)H:Quinone Oxidoreductases. Microbiology and Molecular Biology Reviews 68, 603–616 (2004).

56. H. J. Murff, et al., Metformin use and incidence cancer risk: evidence for a selective protective effect against liver cancer. Cancer Causes Control 29, 823–832 (2018).

57. Y. Kita, et al., Metformin Prevents and Reverses Inflammation in a Non-Diabetic Mouse Model of Nonalcoholic Steatohepatitis. PLoS ONE 7, e43056 (2012).

